# Bacterial species and surface structures shape gene transfer and the transcriptional landscape during active conjugation

**DOI:** 10.1101/2025.06.12.659224

**Authors:** Galain C Williams, Jillian C Danne, Simon Crawford, Jihane Homman-Ludiye, Xenia Kostoulias, Bailey Heron, Tasnia Islam, Melvin Yong, Yunn-Hwen Gan, Yogitha N Srikhanta, Dena Lyras

## Abstract

Conjugative plasmids drive bacterial evolution and niche adaptation, yet how their active acquisition reshapes host transcription remains poorly understood. Most studies focus on stable plasmid carriage, overlooking the dynamic transcriptional changes during conjugation itself. Here, we show that active RP4 conjugation triggers an immediate, host- and surface factor-dependent transcriptional response. This includes activation of non-SOS stress pathways, motility, exopolysaccharide production, anaerobic respiration, and metabolic adaptation. These responses do not inhibit conjugation, suggesting they serve to maintain host homeostasis. Capsule expression blocks these responses by preventing conjugation, and also inhibits RP4 activation by physically blocking donor-recipient contact. Unexpectedly, RP4 overcomes this barrier by exploiting single-cell variation in recipient capsule thickness, successfully conjugating with thin-capsulated recipients. These findings reveal a striking interplay between plasmid, host, and surface architecture in shaping the conjugation transcriptional landscape, with broad implications for plasmid dissemination and bacterial evolution.

## Introduction

Conjugative plasmids are key drivers of horizontal gene transfer, enabling rapid trait acquisition in bacteria (1). Critically, their spread through bacterial populations is closely linked to the rise of antibiotic resistance across clinical and environmental settings (2, 3). Conjugation, the primary mechanism of plasmid dissemination, involves the transfer of single-stranded plasmid DNA from a donor into a neighbouring recipient *via* a type IV secretion system (4). Although traditionally viewed as a donor-driven process, recent seminal studies have revealed that recipients play active, multifaceted roles in modulating conjugation (5–8). For example, the capsule of *Klebsiella pneumoniae* inhibits transfer of many plasmid types (8, 9), whilst *K. pneumoniae* outer membrane proteins act as receptors that selectively engage cognate plasmid-bearing donors (7). Additionally, plasmid carriage induces widespread changes in recipient chromosomal gene expression (10), altering cellular functions beyond the plasmid’s genetic content. These findings highlight a complex interplay between donor, recipient and plasmid that shapes plasmid dissemination and influences host cell physiology and evolution. In this study, we aimed to further dissect this interplay during active conjugation of the broad host range plasmid RP4.

Stable plasmid carriage is known to broadly reprogram host chromosomal gene expression, affecting metabolic, respiratory, transport, virulence, and other pathways (10–14). However, the transcriptional response of recipient cells during active conjugation, particularly involving broad host range plasmids like RP4, remains poorly understood. While many plasmids trigger an SOS response upon entry (15), RP4 does not strongly activate this pathway (15), suggesting alternative mechanisms may be at play. Despite RP4’s widespread use in inter-species gene transfer, its transcriptional impact across diverse species is surprisingly underexplored. For example, RP4 has been shown to supress ammonia oxidation in environmental bacteria (16), but most studies have been conducted in the presence of external compounds such artificial sweeteners (17), or biochar (18), confounding the effects of RP4 acquisition with those of the compounds themselves.

Research on other broad host range plasmids suggests that initial plasmid-host interactions can shape the outcome of transfer. In *Pseudomonas putida*, for instance, R388 transfer induces expression of the *hsdRMS* restriction system, which is counteracted by the R388 anti-restriction protein ArdC (19). Whether RP4 elicits a similar recipient-specific response, or encounters barriers to conjugation, remains unknown. Understanding how recipient species modulate RP4 conjugation is critical, as broad host range plasmids are major vectors in the spread of antibiotic resistance (20).

Beyond transcriptional responses, recipient bacteria also influence conjugation through their cell surface structures. Capsules, thick outer layers of polysaccharide, are well-established barriers to conjugation (8). Paradoxically, they are also associated with increased rates of horizontal gene transfer (21), suggesting a complex and unresolved relationship between capsule presence and gene exchange. This paradox remains poorly understood, in part because capsule-mediated inhibition of conjugation has only been demonstrated in *Klebsiella* species, and the underlying mechanisms are yet to be elucidated.

Using an innovative liquid displacement method to infer capsule volume (8), Haudiquet *et al* showed that larger capsule serotypes more strongly inhibit plasmid transfer, implicating capsule length as a key factor. Given that cell-to-cell contact is essential for RP4 conjugation (22), a plausible, though unproven, hypothesis is that capsules physically obstruct donor-recipient contact, thereby preventing conjugation. To deepen our understanding of how recipient bacteria respond to and modulate broad host range plasmid conjugation, we investigated the transcriptional and structural impact of active RP4 transfer in *Escherichia coli* and *K. pneumoniae*. Both species exhibited distinct transcriptional responses involving diverse host functions including anaerobic respiration, non-SOS stress adaptation, and substrate transport. Notably, the *K. pneumoniae* response was only observed in the absence of capsule, revealing a previously unrecognised role for the capsule in suppressing host transcriptional reprograming during conjugation.

RP4 gene expression was also influenced by the recipient species and was inhibited by the presence of capsule. We confirmed that capsule blocks conjugation across diverse recipients by preventing donor-recipient cell-to-cell contact. Surprisingly, RP4 could exploit single cell variation in capsular thickness to facilitate transfer, bypassing the need for a capsular serotype switch. These findings provide a unified framework for understanding how recipient species and their surface structures shape the transcriptional landscape and dissemination of broad host range plasmids.

## Results

### Active RP4 conjugation elicits triggers non-SOS stress, respiratory and metabolic reprogramming

To map the transcriptional landscape of RP4 conjugation, we first distinguished active conjugation from transconjugant vertical inheritance using a time-course assay (Fig 1a). At 15 minutes, new transconjugants were actively generated without changes in recipient population size, confirming this as a window of active conjugation and ruling out vertical inheritance as a confounding factor. We then performed RNA-seq on a conjugating mixture of laboratory donor *E. coli* LT101(pVS520) and a genetically distant clinical recipient, *E. coli* DLL7555. For comparison, RNA was also extracted from mock conjugations lacking pVS520, and from donor monocultures. The >78,000 core genome differences between DLL7555 and LT101 enabled accurate, strain-specific read assignment using a concatenated LT101-DLL7555 reference genome.

**Figure 1.**
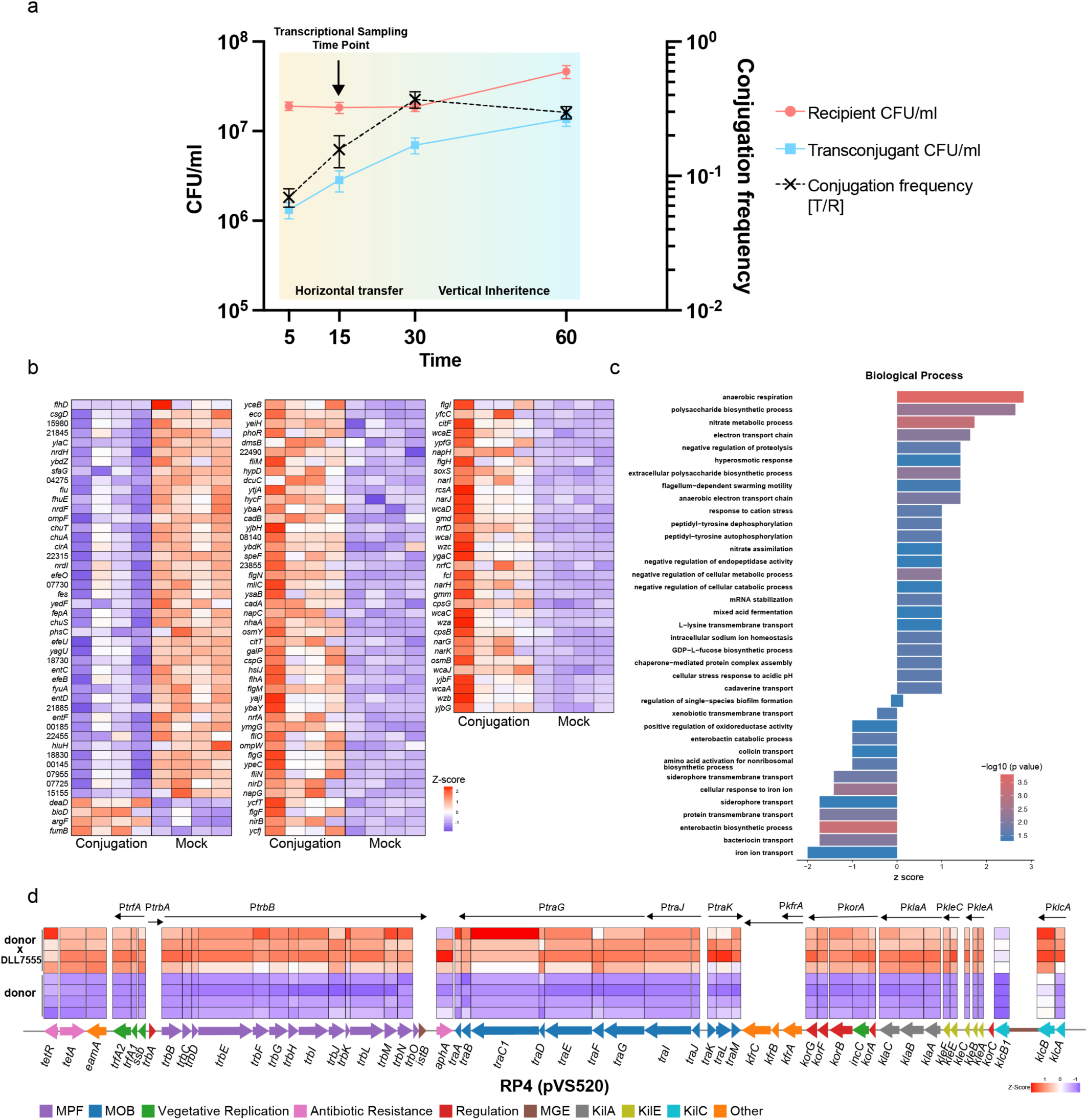
The transcriptional response of an *E. coli* recipient and plasmid RP4 during active conjugation. **a)** *E. coli* DLL7555 conjugation frequency over time shows that RP4 conjugation is active at 15 minutes and completed by 30 minutes. Arrow indicates time for transcriptional sampling. **b)** Differentially expressed DLL7555 genes during active RP4 conjugation. **c)** GO term analysis of enriched genes with the R TopGo package with the weight_01 algorithm. **d)** Differential expression of RP4-encoded genes during active conjugation with *E. coli* DLL7555. Numbers in *E. coli* gene list represent locus tags trimmed of the prefix ACPEGL. Error bars = mean ± SD, N = 4.

RP4 conjugation triggered differential expression of 130 genes in the *E. coli* recipient (Fig 1b), including those involved in adaptation to external stressors such as temperature shock (*cspG*, *hslJ, deaD*), pH regulation (*yagU, ygaC, ybay, speF*), osmotic stress (*osmB, osmY, nhaA*), and other stress-related pathways (*mliC, soxS, rcsA, phoR*). This was accompanied by upregulation of genes involved in colanic acid biosynthesis (*wzc, wzb, cpsB, wcaJ, wcaI, wcaE, wcaC, wcaA, gmd, fcl, wza, wcaD*), exopolysaccharide production (*yjbG, yjbF, yjbH*), and flagella assembly (*fliN, fliM, flgG, flgF, glgI, flgH, flhA, fliO*).

Although conjugative plasmids are known to promote biofilm formation (23), we observed decreased expression of the biofilm master regulator *csgD*. Consistently, co-culture of DLL7555 with LT101(pVS520) resulted in a slight but significant reduction in biofilm formation compared to both LT101 co-culture or monoculture conditions (Supplementary Fig 1).

RP4 conjugation also induced upregulation of genes involved in anaerobic respiration and energy generation (*fdnI, dmsB, napC, narG, narH, narI, hycF, nrfA, nrfD, nrfC, nirB, fumB, dcuC*), overlapping with pathways linked to inorganic nutrient metabolism and nitrogen cycling (*napG, napH, narG, narH, narI, nrfA, narJ, nirB, nirD, narK, napC*). In contrast, genes associated with siderophore, iron transport, and iron homeostasis were downregulated, including *fiu, cirA, fepA, fyuA, efeO, efeU, efeB, fhuE, entD, entC, entF, chuA, chuS, chuT*.

Gene Ontology (GO) term analysis reflected this multifaceted recipient response, with significant enrichment in categories related to stress adaptation, respiratory activity, and metabolic reprogramming (Fig 1c).

By comparing the conjugation condition to the donor alone, we aimed to uncover the global expression profile of RP4 across the entire conjugating population, recognizing that this expression could originate from either donor or transconjugant cells. This analysis revealed a striking upregulation of 47 genes encoded by pVS520, spanning key functional categories including DNA mobilisation (MOB), mating pair formation (MPF), regulation and replication, host-kill systems (*kil*), and other cellular processes (Fig 1d). These findings confirm prior transcriptional studies of RP4 during *E. coli* conjugation (24), demonstrating that that RP4 undergoes intense transcriptional activation during conjugation with this species, with nearly all transcriptional units switched on.

### Capsule shields the recipient from transcriptional reprogramming during conjugation

To assess whether the recipient response to RP4 conjugation was conserved, we repeated our experiment using *K. pneumoniae* B5055 as the recipient. This strain’s hypermucoid capsule also allowed us to investigate how recipient cell surface structures influence the transcriptional landscape during conjugation. Capsules are known barriers to conjugation (8), and prior to RNA-seq, we confirmed that the capsule of *K. pneumoniae* B5055 significantly inhibited RP4 conjugation (Supplementary Fig 2). Since this effect has only been demonstrated in *Klebsiella spp in vitro*, we extended our analysis to *Acinetobacter baumannii* AB5075 and *Pasteurella multocida* VP161, confirming that capsule similarly impedes conjugation in these species (Supplementary Fig 2). This represents the first *in vitro* evidence that capsules broadly inhibit conjugation across multiple genera beyond *Klebsiella*.

To investigate how the capsule influences the transcriptional response to conjugation, we compared gene expression profiles in wild-type and acapsular *K. pneumoniae* recipients. In the wild-type strain, only 10 genes were differentially expressed (including *carB, carA, lysS, clbS, glnH,* 08025*, glnP,* and several uncharacterized loci: 08030, 08050, 08045) (Fig 2a). These genes were associated with pathways such as sucrose catabolism, L-arginine biosynthesis and nucleotide biosynthesis (Fig 2b).

**Figure 2.**
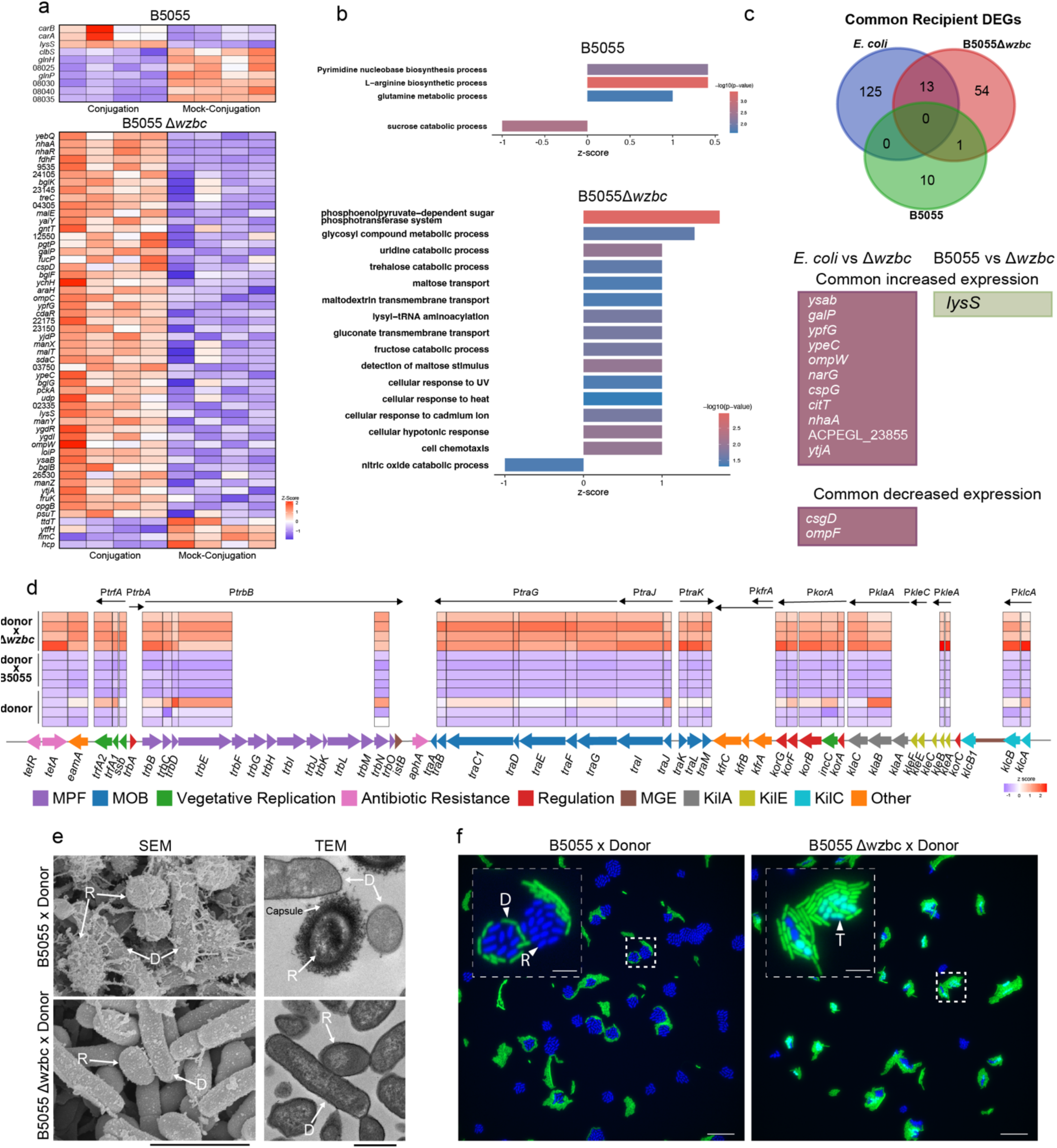
The effect and mechanism of capsule on conjugation and the recipient and plasmid response. **a)** Differentially expressed genes (DEGS) in *K. pneumoniae* B5055 (top) and B5055Δ*wzbc* (bottom) genes during active conjugation or co-culture with an RP4 donor. **b)** GO term analysis of DEG for B5055 (top) and B5055Δ*wzbc*, with raw P-values reported as per TopGO package guidelines for the weight_01 algorithm. **c)** Common DEGS between recipient responses (top), that are increased in expression (middle) or decreased (bottom). **d)** Differentially expressed RP4 genes during active conjugation with B5055Δ*wzbc* (top) B5055 (middle), or the donor alone (bottom). **e)** TEM, and SEM images of conjugation between *K. pneumoniae* B5055 or B5055Δ*wzbc* recipients with *E. coli* LT101. Recipients are distinguished based on the presence of capsule, and shape. **f)** *In situ* visualisation of conjugation between either wildtype or Δ*wzbc* recipients by fluorescent microscopy. All images are representative of ≥2 independent experiments. Abbreviations: D=Donor, R=Recipient, T=Transconjugant. Numbers in *K. pneumoniae* gene lists represent locus tags trimmed of the prefix J6T75_RS. Scale bars: SEM 3 µm, TEM 1 µm, Fluorescence microscopy 20 µm, inset 5 µm.

However, when the acapsular mutant B5055Δ*wzbc* was used as the recipient, the transcriptional response expanded dramatically to 54 genes. This striking increase demonstrates that the capsule not only acts as a physical barrier to conjugation but also shields the recipient from widespread transcriptional reprogramming. By limiting conjugation frequency, the capsule effectively insulates the host transcriptome from the impact of plasmid acquisition.

Building on this observation, we further examined this transcriptional landscape in the absence of capsule. *K. pneumoniae* B5055Δ*wzbc* exhibited upregulation of stress adaptation genes shared with *E. coli* (e.g. *cspD*, *nhaA*), as well as responses unique to this strain (*loiP*, *ychH, hcp*). We also observed broad changes in genes involved in carbohydrate transport (*araH, fucP, galP, manY, manZ, manX, bglF*), catabolism (*treC, fruK pck, bglB, opbG*), and transmembrane transport (*psuT, sdaC, ompC, ttdT, nhaA, yebQ, malE, gntT, ompW*).

Consistent with the *E. coli* response, genes linked to anaerobic growth (*ompW*, *ttdT*) and anaerobic respiration (*fdnG, fdhF, narG*) were also activated. GO term analysis highlighted enrichment in mixed stress responses and altered carbohydrate transport and metabolism (Fig 2b). Using BLAST, we identified 13 homologous genes shared between the two recipient species (*ysab, galP, ypfG, ypeC, ompW, narG, cspG, citT, nhaA,* ACPEGL_23855*, ytjA*, *csgD* and *ompF*), indicating that approximately 10% of the transcriptional response to RP4 is conserved.

### Capsules block conjugation by preventing donor cell activation and cell-to-cell contact

Capsules are potent inhibitors of bacterial conjugation, widely believed to block essential donor-recipient contact, yet this mechanism has not been directly demonstrated. Before examining their role in physical contact, we first asked whether capsules influence the activation of RP4-encoded genes during conjugation. To address this, we re-analysed our transcriptomic dataset comparing wild-type *K. pneumoniae* and its acapsular mutant B5055Δ*wzbc*. When the recipient was encapsulated, no RP4 genes were activated (Fig 2d). In contrast, conjugation with the acapsular mutant triggered expression of 27 genes. Notably, this was fewer than the number activated during conjugation with *E. coli*, suggesting that RP4 gene expression is shaped by the specific recipient species.

To determine whether RP4 gene activation originated from the donor or transconjugant cells, we employed a donor-specific fluorescent reporter assay targeting the relaxase promoter. By using rifampicin to selectively silence recipient transcription, taking advantage of donor rifampicin resistance, we confirmed that relaxase operon activation occurred exclusively in donor cells. Crucially, this activation was blocked when the recipient was encapsulated (Supplementary Fig 3).

These findings reveal that the capsule prevents the donor from sensing the recipient, thereby suppressing transcriptional activation of RP4. This highlights a previously unrecognized role of the capsule in modulating donor behavior during conjugation.

To uncover the mechanism by which capsule inhibits conjugation, we performed imaging studies to directly visualize donor-recipient interactions in the presence and absence of capsule. Transmission and scanning electron microscopy revealed that the capsule of *K. pneumoniae* physically obstructs direct outer membrane contact between donor and recipient cells (Fig 2e, top panels). In contrast, when the capsule was absent, intimate membrane-to-membrane contact was readily observed (Fig 2e, bottom panels), consistent with conditions permissive for conjugation. Unexpectedly, these imaging studies also revealed substantial heterogeneity in capsule thickness and architecture at the single cell level, a phenomenon explored further in a subsequent section (See Fig 3).

**Figure 3.**
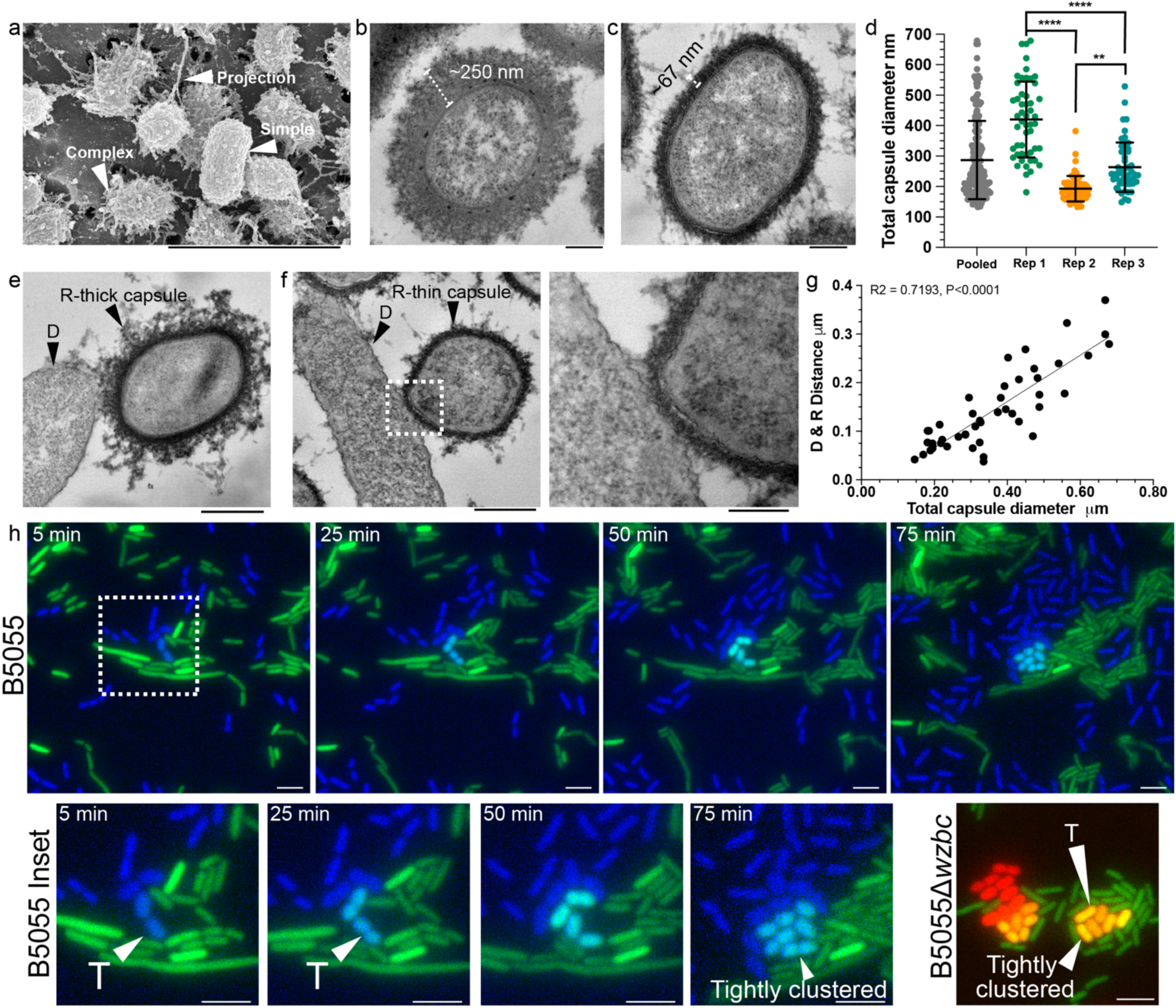
Recipient *K. pneumoniae* capsule thickness variation impacts donor-recipient interactions and conjugation. **a)** SEM imaging of capsular variation and projections. **b-c)** TEM imaging of capsule thickness variation between individual cells. **d)** Variation in capsule thickness within, and between biological replicates (n = ≥50 cells per sample, one-way ANOVA with sidak’s multiple comparison testing P-values = ** ≤ 0.01, **** ≤ 0.0001.). **e-f)** Effect of capsule thickness variation on donor-recipient interactions. **g)** Linear regression analysis of distance between outer membrane of donor and recipient and total capsular diameter (R^2^=.72, N=47, F-test P≤0.0001). **h)** Live cell imaging of B5055 transconjugant formation, showing that B5055 transconjugants arise from recipients intimately contacting the donor and morphologically resemble acapsular B5055Δ*wzbc*. Error bars = mean ± SD. Scale bars: a = 5 µm, bc = 200 nm, ef = 500 nm, f inset = 200 nm, h = 20 µm, h insert = 5 µm. Images are representative of ≥ 2 independent experiments, except for h which depicts the rare formation of a B5055 transconjugant.

To directly test whether capsule-mediated inhibition of cell-to-cell contact blocks conjugation, we performed fluorescence microscopy on agarose pad conjugation assays (Fig 2f). We used an RP4-mobilisable plasmid encoding GFP to track plasmid transfer into recipient cells labelled with BFP on a non-mobilizable plasmid. In conjugations with acapsular mutant B5055Δ*wzbc*, 70% of donor-contacting recipient cells successfully acquired the plasmid (369 transconjugants from 527 mating pairs; Fig 2f, right panel). In stark contrast, only 0.58% of donor-contacting wild-type recipient cells formed transconjugants (2 of 345 mating pairs), and overall mating pair formation was reduced due to capsule-mediated inhibition of direct contact (Fig 2f, left panel). Notably, wild-type *K. pneumoniae* B5055 cells also showed limited contact with each other, indicating that the capsule restricts both intracellular and intercellular interactions.

Together, these results demonstrate that the capsule prevents donor recognition of the recipient by physically blocking cell-to-cell contact, thereby suppressing conjugation.

### RP4 overcomes capsule-mediated conjugation barriers by exploiting capsule thickness and expression variability

During earlier imaging experiments, we observed striking variability in *K. pneumoniae* capsular thickness at the single-cell level, prompting a more detailed analysis using SEM and TEM (Fig 3).

SEM revealed that the capsule forms a complex cell surface matrix, often connecting adjacent cells *via* thick, filamentous projections ranging from 0.5 µm to 4 µm in length. These structures were present in 86% of cells examined (581/676), while a minority displayed smooth, unstructured surfaces (Fig 3a).

TEM further showed that the capsule appears as a diffuse, often non-uniform layer, with substantial variation in thickness between individual cells (Fig 3b-c). Quantification of total capsule diameter across biological replicates revealed a wide range, from 132.8 nm to 678.8 nm, with a mean of 287 nm (Fig 3d). Surprisingly, even under identical growth and processing conditions, capsule thickness varied significantly between replicates, with mean values of 420.2 nm, 192.7 nm, and 263.3 nm, respectively (Fig 3d).

These findings demonstrate that *K. pneumoniae* exhibits pronounced phenotypic heterogeneity in capsule architecture, even within clonal populations, highlighting a potential vulnerability that RP4 may exploit to bypass capsule-mediated conjugation barriers.

To determine whether variability in recipient capsule thickness enables rare conjugation events in wild-type *K. pneumoniae*, we investigated whether thinner-capsule cells were responsible for the small number of transconjugants observed. Since capsule blocks conjugation by preventing close donor-recipient contact (Fig 2g-h), we hypothesized that cells with thinner capsules might permit tighter membrane interactions, allowing plasmid transfer to occur. To test this, we conducted further experiments to directly link capsule heterogeneity with conjugation efficiency.

Supporting this hypothesis, both TEM and SEM imaging revealed that B5055 recipient cells with thinner capsules formed significantly closer associations with donor cells compared to those with thicker capsules (Fig 3 e-f). Quantitative analysis confirmed a strong correlation between capsule thickness and the physical distance separating donor and recipient outer membranes (Fig 3g, R2=0.72, N=47, F test P<0.0001), indicating that thicker capsules physically increase intercellular spacing and reduce the likelihood of conjugation.

Crucially, live-cell fluorescence microscopy showed that transconjugants arose specifically from recipient cells capable of forming intimate contact with donors (Fig 3h). These transconjugant cells appeared in tightly packed clusters and displayed a morphology strikingly similar to the acapsular B5055Δ*wzbc* mutants (Fig 3h, inset), consistent with them being thin-capsulated.

Together, these findings demonstrate that natural heterogeneity in capsule thickness enables a subpopulation of recipient cells to bypass the capsule barrier, facilitating rare but successful conjugation events.

Given the observed variability in capsule thickness across identically treated replicates (Fig 3d), we hypothesized that this heterogeneity could occasionally permit higher conjugation frequencies in the wild-type B5055 *K. pneumoniae* recipient. To test this, we conducted 19 independent conjugation experiments, anticipating that natural fluctuations in capsule thickness between replicates would influence conjugation efficiency. In 4 of these experiments, we observed a markedly elevated mean transfer frequency of 4.2×10^-2^ (Fig 4A), approximately ∼50-fold higher than the median across all replicates, yet still 20-fold lower than that observed with the acapsular B5055Δ*wzbc* mutant (Fig 4a). This suggests that while capsule was still present, its barrier function was partially compromised. Importantly, the resulting transconjugants remained mucoid and string-test positive, confirming they were not spontaneous capsule-deficient mutants.

**Figure 4.**
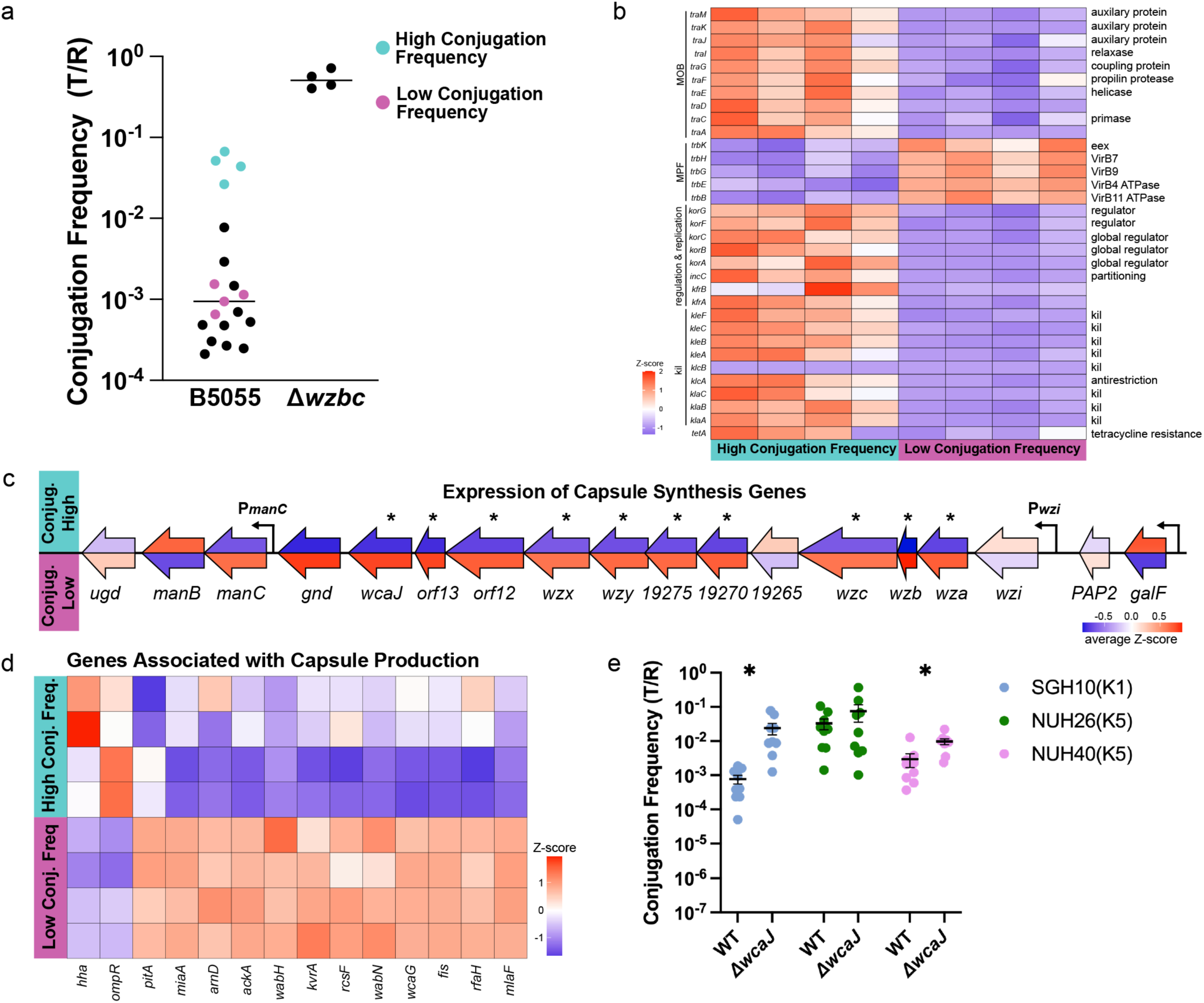
Population level capsule variation enables RP4 conjugation in capsulated *K. pneumoniae*. **a)** Variation in conjugation frequency of *K. pneumoniae* B5055. The conjugation frequency of B5055Δ*wzbc* is shown for context. **b)** Differential expression of RP4 encoded genes between conjugation high and low samples. **c)** Differential expression of genes in the capsule synthesis locus between conjugation high (top) and low (bottom) samples. **d)** Genes associated with capsule production between samples with high or low conjugation frequencies. **e)** The conjugation frequency of RP4 to hypermucoid K1 or K5 *K. pneumoniae* (N=9, P>.05 Mann-Whitney U test). Data are presented as the median (**a**), or the mean+SEM (**d**).

These findings indicate that stochastic variation in capsule expression can transiently weaken its barrier function, enabling rare but enhanced conjugation events in otherwise encapsulated populations.

To investigate the molecular basis of elevated conjugation in wild-type *K. pneumoniae* replicates, we performed RNA-seq to compare RP4 conjugation events and recipient capsule biosynthesis activity between high-frequency and low-frequency conjugation events. In high frequency conjugations, RP4 gene expression resembled that observed in the acapsular B5055Δ*wzbc* mutant (See Fig 2d), characterized by strong upregulation of DNA mobilization genes, but not mating pair formation genes (Fig 4b).

Strikingly, these samples also showed markedly reduced expression of key capsule biosynthesis genes under the control of the P*wzi* promoter (Fig 4c). Beyond *wzi*,13 additional capsule-associated genes (25) were differentially expressed, including the master capsule synthesis regulator *kvrA* (26), and a suite of regulatory and structural genes, such as *hha, ompR, pitA, miaA, arnD, ackA, wabH, rcsF, wabN, wcaG, fis, rfaH* and *mlaF* (Fig 4d). Many other genes with diverse function were also differentially expressed, suggesting a broader physiological shift in these high-frequency conjugation conditions.

Together, these data reveal that reduced capsule gene expression underlies the increased conjugation efficiency in certain wild-type replicates, reinforcing the role of capsule as a dynamic and regulatable barrier to horizontal gene transfer.

Finally, to assess how variation in capsule serotype influences conjugation, we examined the impact of different hypermucoid capsule types on RP4 transfer efficiency. Conjugation with a hypermucoid K1 recipient resulted in a dramatic 31-fold reduction in transfer frequency compared to its acapsular counterpart. In contrast, two hypermucoid K5 recipients showed more modest reductions of 3.3 and 2.2-fold, relative to their respective wild-type controls. These findings demonstrate that while capsule consistently acts as a barrier to conjugation, the magnitude of its inhibitory effect varies by serotype and is independent of if the serotype is hypermucoid. This highlights that both dynamic capsular thickness and serotype-specific properties govern conjugation efficiency through mechanisms that are partially independent of capsular serotype.

## Discussion

We investigated how recipient species and surface structures shape the early transcriptional response and ultimate success of broad host range plasmid conjugation. Our transcriptomic analyses revealed that plasmid acquisition is a highly transcriptionally active process, triggering non-SOS stress responses and metabolic reprogramming in recipient cells. Notably, we found that the bacterial capsule acts as a protective barrier, shielding recipients from this transcriptional upheaval by physically blocking donor-recipient contact and thereby preventing donor activation.

We further demonstrated that this inhibitory effect is not unique to *K*. *pneumoniae* since capsules from *A. baumannii and P. multocida* also suppressed conjugation, establishing capsules as broad-spectrum barriers to horizontal gene transfer. Strikingly, we discovered that the capsule is not an absolute barrier. RP4 can exploit natural heterogeneity in capsule thickness at the single-cell level to bypass this defense, enabling sporadic but successful plasmid transfer into otherwise resistant populations.

Together, our findings provide a unified framework for understanding how recipient identity and surface structure variability govern the transcriptional landscape and efficiency of conjugation. This work offers new insights into how broad host range plasmids overcome physical barriers to disseminate across diverse bacterial species.

A key finding of this study is that conjugation induces a far more extensive transcriptional response than previously appreciated, extending well beyond the canonical SOS response (27). Across both *E. coli* and *K. pneumoniae*, conjugation triggered broad non-SOS stress responses and widespread reprogramming of respiratory, metabolic, substrate transport, and other pathways. This likely reflects the substantial energetic and homeostatic burden associated with establishing a new plasmid, consistent with the commonly observed growth defects in newly formed RP4 transconjugants (28).

Despite species-specific differences, approximately 10% of the recipient transcriptional response was shared between *E. coli* and *K. pneumoniae*. Notably, two conserved genes, *ompW* and *nhaA*, have previously been shown to influence the conjugation efficiency of multiple plasmids, including R388 and F-like plasmids (7, 29). OmpW, typically expressed during the transition to anaerobic respiration (30), is targeted by mating pair stabilization proteins during conjugation (7). NhaA, a key regulator of sodium ion, pH, and osmotic balance (31), also contributes to cell envelope integrity (32), suggesting a protective role against conjugation-induced envelope stress.

Importantly, this transcriptional upheaval was largely absent in wild-type *K*. *pneumoniae*, where capsule production effectively suppressed the recipient response. Although only a small number of genes were differentially expressed in the encapsulated strain, GO-term analysis still identified numerous redundant terms, a known limitation of this method (33). In contrast, the broader and more diverse gene expression changes observed in *E. coli* and the acapsular mutant yielded a wide array of non-redundant GO terms, reinforcing the conclusion that capsule acts as a transcriptional shield during conjugation.

Capsules have long been implicated as barriers to conjugation (9, 34), but here we provide the first direct visual evidence that they physically block the essential donor-recipient contact required for mating pair formation. Our findings reveal that this physical barrier has two critical consequences for conjugation. First, by preventing plasmid transfer, the capsule acts as a protective shield for the recipient, insulating it from the extensive transcriptional reprogramming and stress responses triggered by plasmid acquisition. We propose that this shielding effect may confer a selective advantage in competitive environments, where maintaining cellular homeostasis is crucial for survival.

Second, we show that the capsule effectively masks the recipient from the donor, preventing the donor from initiating conjugation. Although the necessity of direct cell-to-cell contact for conjugation has been recognized since the 1950s (35), the role of the recipient during this contact has remained poorly defined. In the case of RP4, stable donor-recipient contact is required for conjugative junction formation (36), yet previous genetic screens in *E. coli* have failed to identify recipient genes essential for RP4-mediated transfer (37).

By demonstrating that contact with a capsulated recipient is insufficient to trigger donor activation, we provide compelling evidence that a recipient-encoded envelope factor, obscured by the capsule, is required to initiate donor transcriptional activation. This aligns with recent findings that R388 pilus expression is contact-dependent (38), but our data suggest that, for RP4, it is the formation of the relaxosome, rather than pilus expression, that is contingent on recipient recognition. Further, we show that contact with capsule is not sufficient to trigger donor activation, showing that contact alone is not sufficient for conjugation.

Unexpectedly, we found that the capsule is not an absolute barrier to conjugation, but rather a leaky one, where natural variation in capsule thickness at the single-cell level permits occasional RP4 transfer. Previous studies had indirectly linked capsule volume to the degree of conjugation inhibition (8), but here we provide direct evidence that capsule thickness precisely determines the physical distance between donor and recipient membranes. This spatial separation in critical, as it governs the likelihood of successful mating pair formation.

Capsular heterogeneity is well-documented across many bacterial species (39–42), and our findings reveal a novel functional consequence of this variability: it modulates horizontal gene transfer. In *K. pneumoniae*, capsule expression is regulated by complex networks involving over 70 genes (25, 43) and is largely non-phase-variable (25, 44, 45). Our data suggest that the observed variation in capsular thickness is predominantly stochastic, with no consistent pattern of expression across the population.

While the functional significance of single-cell heterogeneity in conjugation has only recently begun to emerge (46, 47), our study uncovers a new dimension, specifically, that dynamic, stochastic variation in recipient capsule thickness enables broad host range plasmids like RP4 to breach an otherwise effective barrier and disseminate across encapsulated bacterial populations.

In conclusion, our study reveals that conjugation is a transcriptionally demanding process for recipient cells, extending well beyond the classical SOS response, and that this response is shaped by both species identity and surface architecture. We demonstrate that the bacterial capsule plays a dual role in modulating conjugation: it physically blocks donor-recipient contact, thereby preventing donor activation, and it shields the recipient from the transcriptional stress associated with plasmid acquisition. RP4 can exploit natural, single-cell variation in capsule thickness to bypass this defense and successfully transfer into encapsulated populations.

These findings have broad implications for understanding how conjugative plasmids, carrying antibiotic resistance, virulence factors, and metabolic traits, navigate physical barriers to spread across diverse bacterial species. Capsule heterogeneity emerges as a critical factor influencing the success of horizontal gene transfer, with important consequences for microbial evolution and the dissemination of clinically relevant traits.

## Materials and Methods

### Bacterial strains, plasmids, and growth conditions

Table 1 lists the bacterial strains and plasmids used in this work. For routine growth, *E. coli* and *K. pneumoniae* strains were cultured in low-salt LB at 37^°^C shaking in liquid at 200 rpm or on solid media. Media was supplemented with antibiotics where necessary: Ampicillin (100 µg ml^-1^), rifampicin (150 µg ml^-1^), erythromycin (150 µg/ml), spectinomycin (50 µg/ml) and tetracycline (10 µg ml^-1^).

**Table 1.**
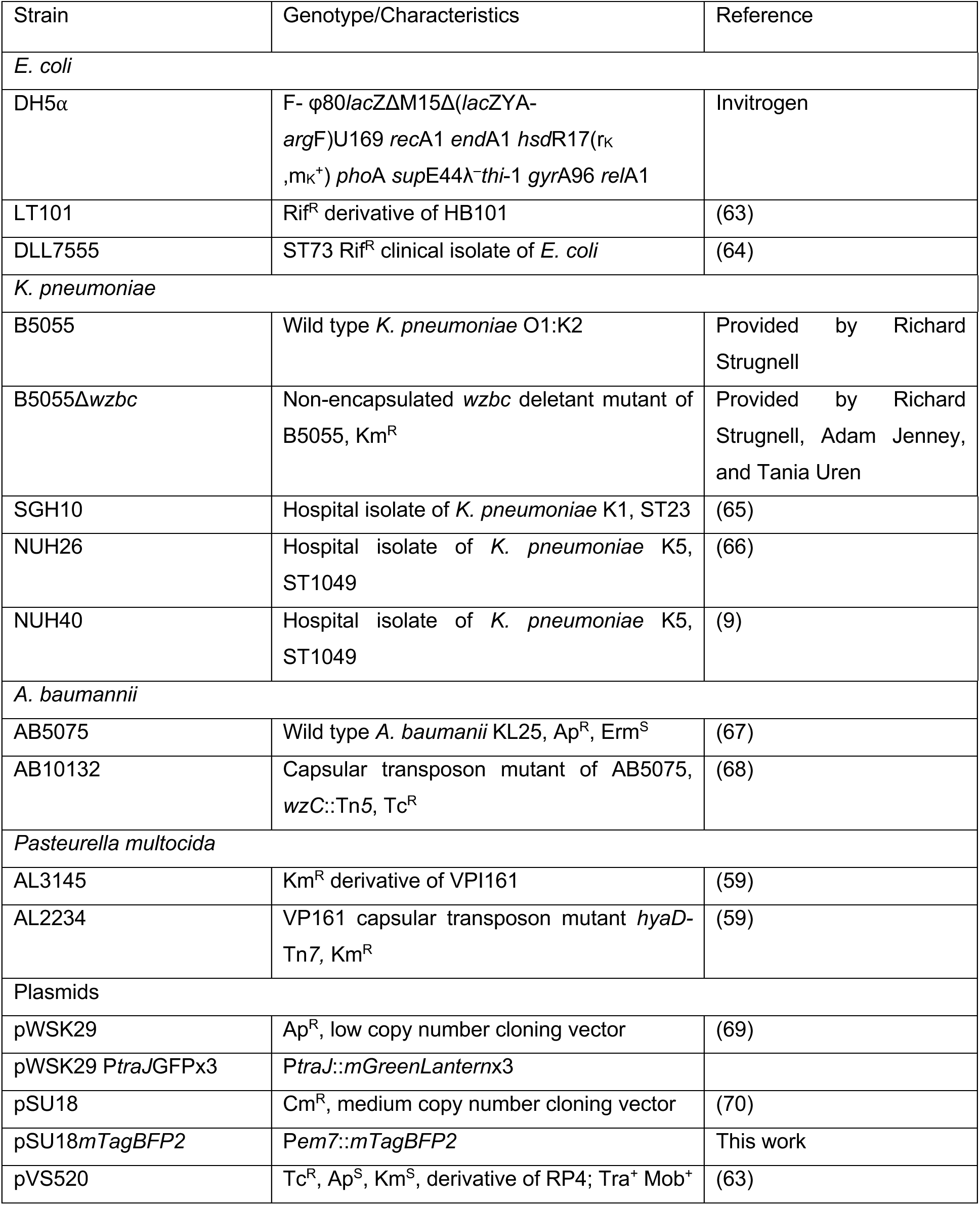
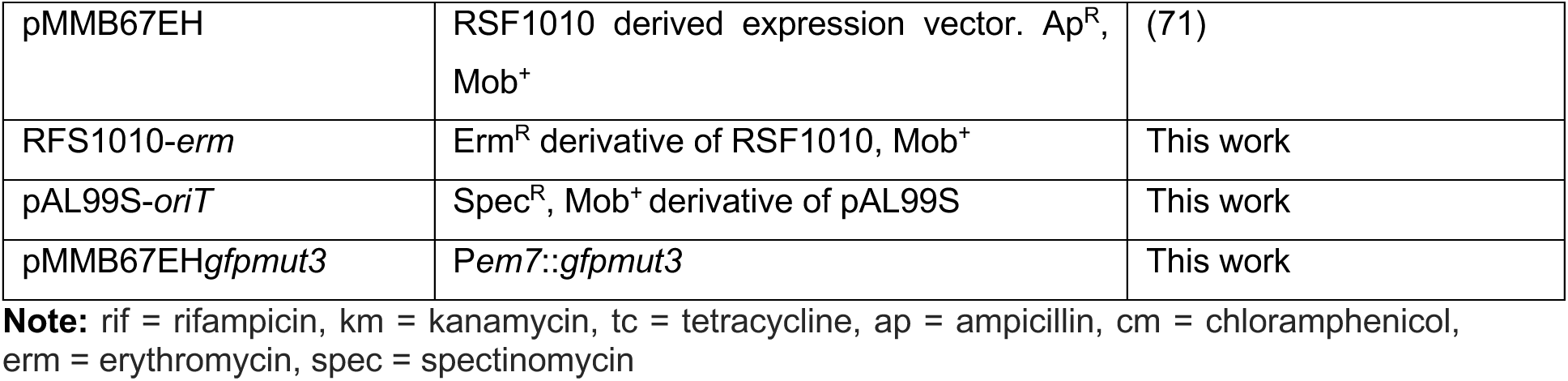
Bacterial strains and plasmids used in this work.

### Strain and plasmid construction

Prior to use as recipients, wildtype and Δ*wzbc K. pneumoniae*, along with *E. coli* DLL7555, were electroporated with the low copy number vector pWSK29 to provide an ampicillin counter selection maker for conjugation. Fluorescent reporter plasmids were made as follows. *GFPmut3* or *mTagBFP2* were fused to a synthetic *em7* promoter (48) and synthesised as gBlocks (IDT). These were cloned into the XbaI and BamHI site of pMMB67EH or pSU18 respectively through restriction digestion cloning. pWSK29P*traJ*GFPx3 was similarly constructed, except that three translationally coupled copies of *mGreenLantern* were fused to a 367 bp region upstream of the *traJ* start codon. Each *mGreenLantern* subunit was independently codon optimised until each shared < 80% sequence homology.

### Conjugation assays

Overnight liquid cultures of donor and recipient were diluted 1:50 in fresh LB and grown for 4 hours before being washed once in PBS and normalized to an OD_600_ 1.0. 500 µl of donor and recipient culture was passed through a 0.44 µm nitrocellulose membrane filter (Whattman) and incubated on LB agar at 37°C for 15 minutes before extraction in 2 ml of PBS by vigorous vortexing. 15 minutes was chosen as a timepoint to prevent secondary transmission of plasmid RP4, and to minimize growth of the recipient which can skew transfer frequencies. For routine assays, recipient and transconjugant CFU/ml were quantified by plating onto ampicillin, and ampicillin + tetracycline respectively. For *A. baumannii*, transconjugants were quantified on ampicillin + erythromycin using mobilizable plasmid RSF1010 *erm* as a surrogate for RP4 transfer. For *P. multocida,* kanamycin + spectinomycin using mobilizable plasmid pAL99S-*oriT* as a surrogate for RP4 transfer was performed. Conjugation frequency was calculated as the transconjugant CFU/ml divided by recipient CFU/ml.

### RNA extraction and sequencing

Conjugations were fixed after 15 minutes of incubation by vortexing mating filters in RNAprotect (Qiagen). Fixed samples were pelleted at 13,300 RPM for 5 minutes, the RNAprotect decanted and samples stored at −80°C for later processing. RNA was extracted from the DLL7555 samples using an RNeasy RNA extraction kit (Qiagen) as per the manufacturer’s instructions with an on-column DNase treatment (Qiagen). Due to its thick capsule, RNA from *K. pneumoniae* was isolated with hot-phenol as previously described (49) and further purified using an RNeasy kit with on column-DNase treatment.

RNA from DLL7555-related experiments was sequenced at the MHTP Medical Genomics Facility at the Monash Translational Facility. Samples were quality checked by Bioanalyzer (Agilent), and libraries generated using the Illumina stranded total RNA-seq protocol with ribosomal RNA depletion performed using the bacterial Ribo-Zero rRNA removal kit (Illumina). Sequencing was performed on an Illumina NextSeq550 on high output mode with chemistry version 2.5, using the Illumina protocol 15046563 version 2. RNA from *K. pneumoniae* related experiments was sequenced at Micromon Genomics. Samples were quality checked by Fragment Analyzer (AATI) and RNA library construction performed using MGI Tech MGIEasy RNA Directional Library Prep Kit V2 (1000006386). Ribosomal RNA was depleted using the rRNA depletion kit bacteria (NEB). Resulting libraries were sequenced on an MGIEasy MGISEQ2000RS using V2 chemistry. For all experiments, samples were sequenced to a depth that would ensure least 10 million reads per isolate in the sample.

### Transcriptomic analysis

Prior to analysis, we assessed the suitability of our two genetically distant *E. coli* donor and recipient strains for RNA-seq analysis. Genetic variation between the donor and recipient was assessed using snippy version 4.6 (50) which detected >75,000 genetic variants including >68,000 SNPs which were equally distributed across the recipient genome (Supplementary Fig 4). This number was in line with previous single species co-culture RNA-seq experiments (51). Although the two strains were sufficiently genetically distinct, 53 genes that were identical between DLL7555 and HB101 were excluded from the final analysis. RNA reads were aligned against a concatenated sequence of the donor (accession, CP184671.1), recipient (DLL7555, accession CP186931.1 or B5055, assembly GCF_024138975.1), and pVS520 sequences (CP184672.1). The sequence for HB101 (Rif^S^ version of LT101) was used as the donor sequence owing to its availability. Read alignment was performed with subread version v1.5.1 (52) against the concatenated sequence which discarded reads of ambiguous donor/recipient origin. >99.8% of donor originating reads were filtered by this strategy for both B5055 and DLL755 matings, which was assessed by alignment of RNA isolated from the donor only condition against the concatenated sequence. Read counting and assignment were performed via FeatureCounts (53). Differential gene expression was analysed with the Voom/Limma method in Degust v4.1 (https://drpowell.github.io/degust/) with a minimum of 3 counts per million reads across 3 samples. At this stage, an additional 5 genes were subsequently excluded from the DLL7555 analysis due to the detection of donor originating reads which were not successfully filtered earlier. Genes were considered differentially expressed if they had a Log2 fold change ≥ 0.585, a P value ≤ 0.05, and an FDR ≤ 0.05. GO term analysis was performed using the TopGO R package v2.58 (54), with the weight_01 algorithm using raw P-values due to the structured dependency of GO terms as per the package guidelines. eggNOG mapper v2.1.8 (55) was used to assign GO-terms to the GenBank recipient proteomes. Gene and pathway functions were also assessed using the ecocyc database (56). Heatmaps and GO term plots were generated using ggplot2 v3.5.2 (57) and the geneviewer package version 0.1.1 (https://github.com/nvelden/geneviewer).

### Scanning and Transmission Electron Microscopy

SEM was performed on *K. pneumoniae* cells during conjugation, or in stationary phase when capsule is highly expressed (25). 500 µl of cells were collected onto a 0.4 µm polycarbonate membrane, and the capsular ultrastructure preserved by lysine acetate, ruthenium red and osmium (LRR) fixation (58), followed by graduated dehydration in ethanol, critical point drying, and gold sputter coating as previously described (59). Samples were imaged on a field emission NovaSEM 450 SEM with an acceleration voltage of 10 kV in secondary electron mode with a working distance of ∼4.5mm. Images were taken under immersion mode through the lens detector.

For TEM imaging, cells were collected on polycarbonate membranes and prepared with LRR fixation as for SEM. Fixed membranes were then cut into small rectangles no longer than 1 mm cubed, dehydrated and then infiltrated with Epon resin (60). Each membrane was placed on a glass slide cell-side up, and a BEEM® capsule containing 100% resin positioned over the flat membrane before polymerisation at 60°C for 48 hours. The resin-filled BEEM® capsule and slide were cooled in liquid nitrogen and then immediately separated, leaving the flat membrane with cells in a longitudinal orientation on the surface of the resin block. Cells were ultrathin sectioned using a Leica UC7 ultramicrotome and Diatome diamond knife and then contrasted with uranyl acetate and lead citrate. Cells were imaged on a JEOL JEM-1400 Plus TEM at 80 KeV, equipped with a high sensitivity bottom mount CMOS ‘Flash’ camera. Total capsule diameter was measured from the average of 5 total cell widths from which the cell body width was subtracted, as described previously (61).

### Fluorescent Microscopy and Image analysis

Conjugation was analysed by fluorescent microscopy by inoculation of 1 µl of OD_600_ 0.1 recipient and donor onto LB 1% agarose pads. Pads were incubated in parafilm sealed 25 mm fluorodish chamber (WPI) for 2 hours at 37°C. Small strips of moist KimWipe® were included in the chamber to prevent pad dehydration. Images were acquired using a 63x/1.30 oil objective on a Leica DMI-8 widefield microscope equipped with a Lecia K8 sCMOS monochromatic camera using Lecia LasX software. Mating pair productivity was assessed by counting transconjugants that were in direct contact with a donor and dividing that by the total number of recipients in contact with a donor

### Fluorescent reporter assays

Fluorescent measurements were taken on a Tecan infinite pro 200 plate reader. For measurement of P*traJ*, 15 µl of *K. pneumoniae* was inoculated onto a 200 µl LB 1% agarose pad with rifampicin in a 96-well special optics microtiter plate (Corning). After 5 minutes of incubation and air drying, 15 µl of donor culture was spotted on top and immediately afterwards the OD600 and fluorescent intensity (488 nm / 520 nm) were measured in 5-minute intervals at 37°C. Fluorescent intensity was then normalised to OD600 and analysed as change over time by calculating fold-change against the 0-time point value.

## Author Contributions

G.W. co-designed experiments, performed the majority of the experimental work and co-wrote the manuscript. J.D. and S.C. performed the TEM and assisted with SEM imaging. J.H. assisted with fluorescent microscopy imaging and analysis. X.K. assisted with RNA sequencing. B. H. assisted with fluorescent microscopy. T. I. assisted with conjugation time course experiments. M.Y. and Y.H.G assisted with the *K. pneumoniae* conjugation experiments and experimental design. Y.N.S. co-wrote manuscript & co-designed the majority of experiments. D.L. co-designed all experiments and co-wrote the manuscript.

## Acknowledgments

We would like to acknowledge Dr Francesca Short for providing strains and protocols for RNA isolation, and Dr Tom Smallman and Prof John Boyce for providing strains. The authors acknowledge use of the facilities and the assistance of Alex Fulcher from Monash Micro Imaging. The authors acknowledge the use of instruments and assistance at the Monash Ramaciotti Centre for Cryo-Electron Microscopy, a Node of Microscopy Australia. The authors acknowledge the MHTP Medical Genomics Facility for assistance with RNA sequencing.

## Data availability

Transcriptomic data have been submitted to the NCBI SRA under the bioproject accession PRJNA1286074

---

**Supplementary Figure 1:**
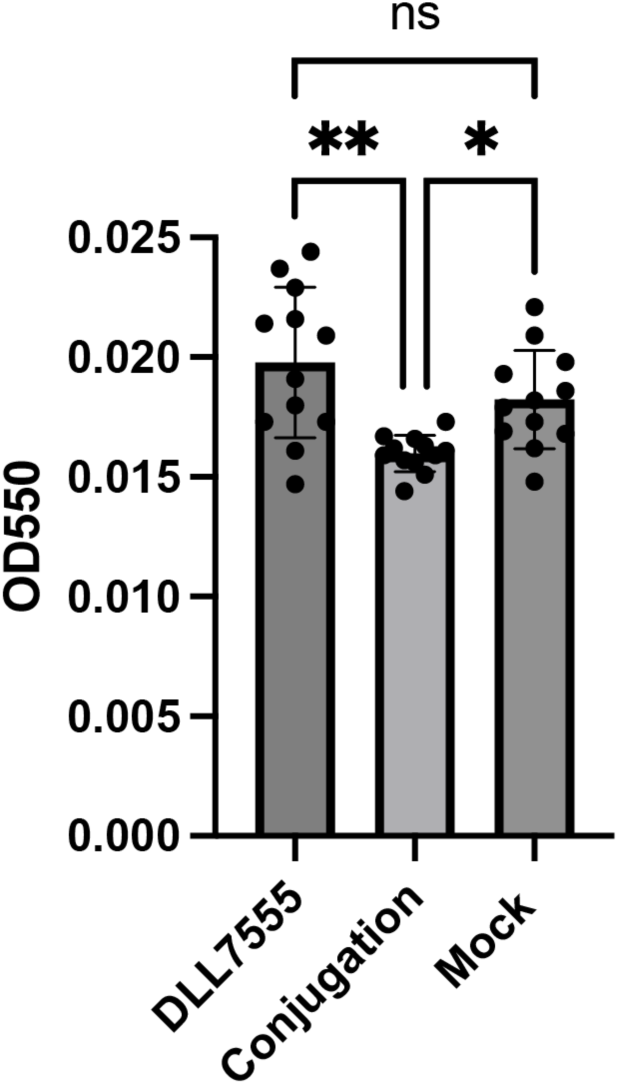
Donor LT101(pVS520) and Recipient DLL7555 co-culture causes a minor reduction in biofilm formation. Biofilms were grown as in (62). Briefly, 1:100 dilutions of overnight broth cultures of donor, recipient and mock donor in fresh LB medium were mixed in 1:1 in a flat bottomed 96 well tray and grown for 24 hours at 37°C. Outer wells were not used to avoid evaporation effects. Following incubation, the liquid portion of the culture was removed and plated to confirm transconjugants were present and therefore conjugation had occurred. The biofilms were washed in water, and quantitated by staining with 0.1% crystal violet, which after washing was solubilized with 30% acetic acid and measured by absorbance reading at 550 nm standardized against 30% acetic acid. Error bars = mean ± SD, N = 4 biological replicates repeated 3 times). Significance testing was performed with a Kruskal Wallis one way ANOVA with Dunn’s multiple comparison testing P-values = *** ≤ 0.01, **** ≤ 0.0001.).

**Supplementary Figure 2:**
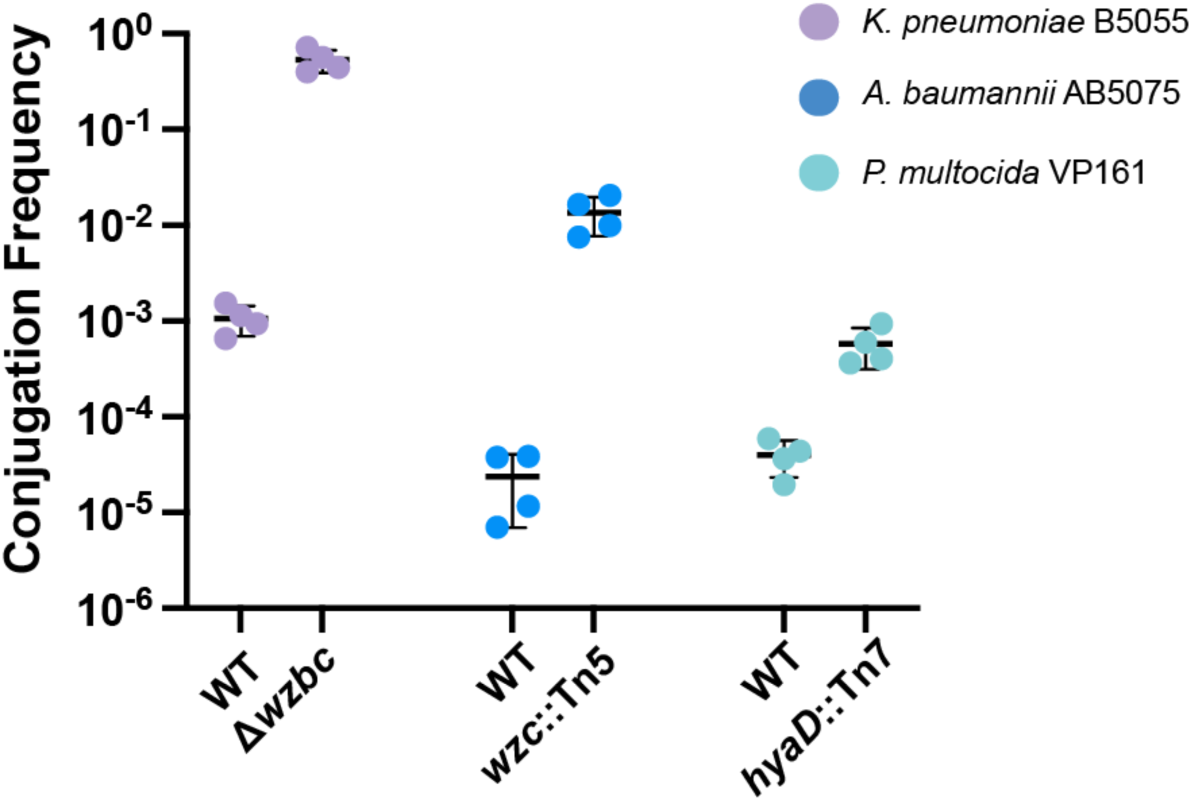
Capsule is a barrier to conjugation in diverse species. The conjugation frequency of RP4 or RP4-mobilizable plasmids from *E. coli* LT101 to K*. pneumoniae* B5055, *A. baumannii* AB5075, or *P. multocida* VP161 and their unencapsulated derivatives (Mean±SD, N=4, P>.05 Mann-Whitney U test).

**Supplementary Figure 3:**
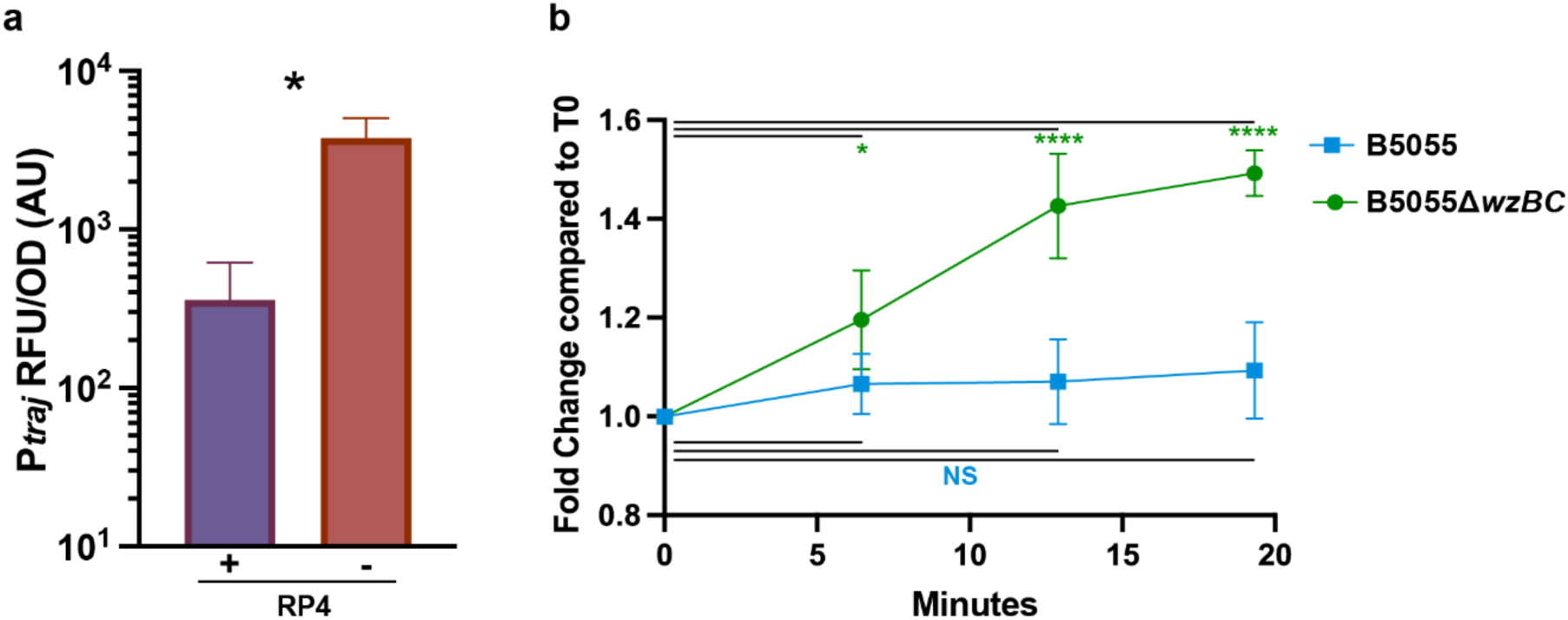
Capsule inhibits activation of the RP4 relaxase promoter P*traJ* within the donor. A fluorescent reporter was constructed for the RP4 relaxase promoter P*traJ* by fusing it to three tandem low sequence-homology GFP reporters. Three tandem GFPs were used because fluorescence from 1 GFP reporter could not be detected, likely because P*traJ* is a weak promoter **a)** The presence of RP4 is known to repress P*traJ* and therefore measurement of GFP intensity in the presence or absence of RP4 confirmed the reporter functioned as expected (N = 4, P<0.05, Mann-Whitney U-Test). **b)** Measurement of P*traJ*x3GFP in the donor cell was achieved by performing conjugations in a 96-well tray on agar containing rifampicin which inhibits transcription only in the recipient cell. Significance testing was performed with a one-way ANOVA with Sidak’s multiple comparison. Error bars = mean ± SD

**Supplementary Figure 4:**
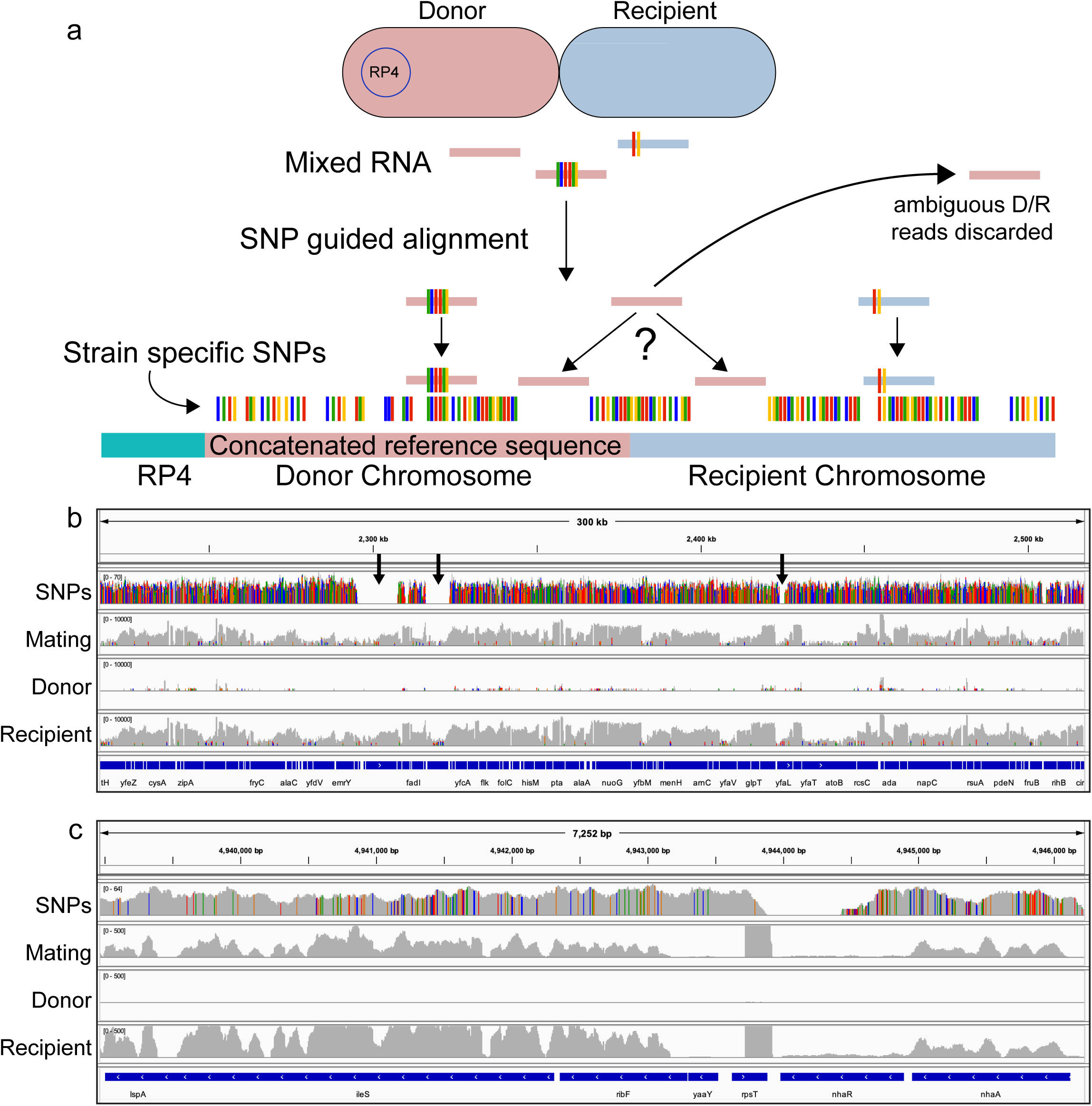
SNP and RNA-seq read alignment for HB101(pVS520) x DLL7555. a) Schematic overview of RNA-seq alignment strategy. b-c) IGV plots depicting the coverage of SNPs, and the alignment of mating, pure recipient and pure donor read coverage against a recipient portion of the concatenated reference sequence at a global (b) and local (c) level. Gaps indicated by black arrows in represent genome segments unique to DLL7555. The reads are log scaled in (b).

## References

1. S. Redondo-Salvo et al., Pathways for horizontal gene transfer in bacteria revealed by a global map of their plasmids. Nat Commun 11, 3602 (2020).

2. S. Castañeda-Barba, E. M. Top, T. Stalder, Plasmids, a molecular cornerstone of antimicrobial resistance in the One Health era. Nature Reviews Microbiology 22, 18–32 (2024).

3. W. Matlock et al., Enterobacterales plasmid sharing amongst human bloodstream infections, livestock, wastewater, and waterway niches in Oxfordshire, UK. Elife 12 (2023).

4. C. Virolle, K. Goldlust, S. Djermoun, S. Bigot, C. Lesterlin, Plasmid Transfer by Conjugation in Gram-Negative Bacteria: From the Cellular to the Community Level. Genes (Basel) 11 (2020).

5. L. Toribio-Celestino, A. San Millan, Plasmid-bacteria associations in the clinical context. Trends in Microbiology 33, 937–947 (2025).

6. M. P. Garcillán-Barcia, F. de la Cruz, E. P. Rocha, The extended mobility of plasmids. Nucleic Acids Research 53, gkaf652 (2025).

7. W. W. Low et al., Mating pair stabilization mediates bacterial conjugation species specificity. Nature Microbiology 7, 1016–1027 (2022).

8. M. Haudiquet et al., Capsules and their traits shape phage susceptibility and plasmid conjugation efficiency. Nature Communications 15, 2032 (2024).

9. M. Yong et al., Dominant Carbapenemase-Encoding Plasmids in Clinical Enterobacterales Isolates and Hypervirulent Klebsiella pneumoniae, Singapore. Emerg Infect Dis 28, 1578–1588 (2022).

10. K. Billane, E. Harrison, D. Cameron, M. A. Brockhurst, Why do plasmids manipulate the expression of bacterial phenotypes? Philosophical Transactions of the Royal Society B: Biological Sciences 377, 20200461 (2022).

11. H. Lee, K. S. Ko, Effect of multiple, compatible plasmids on the fitness of the bacterial host by inducing transcriptional changes. Journal of Antimicrobial Chemotherapy 76, 2528–2537 (2021).

12. C. Huang et al., Comparative Analysis of Transcriptome and Proteome Revealed the Common Metabolic Pathways Induced by Prevalent ESBL Plasmids in Escherichia coli. International Journal of Molecular Sciences 24, 14009 (2023).

13. A. San Millan, M. Toll-Riera, Q. Qi, R. C. MacLean, Interactions between horizontally acquired genes create a fitness cost in Pseudomonas aeruginosa. Nature Communications 6, 6845 (2015).

14. K. Schaufler et al., Carriage of Extended-Spectrum Beta-Lactamase-Plasmids Does Not Reduce Fitness but Enhances Virulence in Some Strains of Pandemic E. coli Lineages. Front Microbiol 7, 336 (2016).

15. Z. Baharoglu, D. Bikard, D. Mazel, Conjugative DNA Transfer Induces the Bacterial SOS Response and Promotes Antibiotic Resistance Development through Integron Activation. PLOS Genetics 6, e1001165 (2010).

16. C. Zhao et al., Horizontal transfer of the multidrug resistance plasmid RP4 inhibits ammonia nitrogen removal dominated by ammonia-oxidizing bacteria. Water Research 217, 118434 (2022).

17. Z. Yu, Y. Wang, J. Lu, P. L. Bond, J. Guo, Nonnutritive sweeteners can promote the dissemination of antibiotic resistance through conjugative gene transfer. Isme j 15, 2117–2130 (2021).

18. Y. Wu et al., Biochar Effectively Inhibits the Horizontal Transfer of Antibiotic Resistance Genes via Restraining the Energy Supply for Conjugative Plasmid Transfer. Environ Sci Technol 56, 12573–12583 (2022).

19. L. González-Montes, I. Del Campo, M. P. Garcillán-Barcia, F. de la Cruz, G. Moncalián, ArdC, a ssDNA-binding protein with a metalloprotease domain, overpasses the recipient hsdRMS restriction system broadening conjugation host range. PLoS Genet 16, e1008750 (2020).

20. C. Coluzzi, E. P. C. Rocha, The Spread of Antibiotic Resistance Is Driven by Plasmids Among the Fastest Evolving and of Broadest Host Range. Molecular Biology and Evolution 42 (2025).

21. O. Rendueles, J. A. M. de Sousa, A. Bernheim, M. Touchon, E. P. C. Rocha, Genetic exchanges are more frequent in bacteria encoding capsules. PLoS Genet 14, e1007862 (2018).

22. T. D. Lawley, G. S. Gordon, A. Wright, D. E. Taylor, Bacterial conjugative transfer: visualization of successful mating pairs and plasmid establishment in live Escherichia coli. Molecular Microbiology 44, 947–956 (2002).

23. J. M. Ghigo, Natural conjugative plasmids induce bacterial biofilm development. Nature 412, 442–445 (2001).

24. M. Miyakoshi, Y. Ohtsubo, Y. Nagata, M. Tsuda, Transcriptome Analysis of Zygotic Induction During Conjugative Transfer of Plasmid RP4. Frontiers in Microbiology Volume 11 - 2020 (2020).

25. M. J. Dorman, T. Feltwell, D. A. Goulding, J. Parkhill, F. L. Short, The Capsule Regulatory Network of Klebsiella pneumoniae Defined by density-TraDISort. mBio 9 (2018).

26. M. Palacios et al., Identification of Two Regulators of Virulence That Are Conserved in Klebsiella pneumoniae Classical and Hypervirulent Strains. mBio 9 (2018).

27. H. D. Curtsinger, S. Martínez-Absalón, Y. Liu, A. J. Lopatkin, The metabolic burden associated with plasmid acquisition: An assessment of the unrecognized benefits to host cells. Bioessays 10.1002/bies.202400164, e2400164 (2024).

28. H. Prensky, A. Gomez-Simmonds, A. C. Uhlemann, A. J. Lopatkin, Conjugation dynamics depend on both the plasmid acquisition cost and the fitness cost. Molecular Systems Biology 17, e9913 (2021).

29. D. Pérez-Mendoza, F. de la Cruz, Escherichia coli genes affecting recipient ability in plasmid conjugation: Are there any? BMC Genomics 10, 71 (2009).

30. M. Xiao, Y. Lai, J. Sun, G. Chen, A. Yan, Transcriptional Regulation of the Outer Membrane Porin Gene ompW Reveals its Physiological Role during the Transition from the Aerobic to the Anaerobic Lifestyle of Escherichia coli. Front Microbiol 7, 799 (2016).

31. T. A. Krulwich, G. Sachs, E. Padan, Molecular aspects of bacterial pH sensing and homeostasis. Nat Rev Microbiol 9, 330–343 (2011).

32. Z. Mao et al., NhaA facilitates the maintenance of bacterial envelope integrity and the evasion of complement attack contributing to extraintestinal pathogenic Escherichia coli virulence. Infect Immun 91, e0003923 (2023).

33. R. Barriot, D. J. Sherman, I. Dutour, How to decide which are the most pertinent overly-represented features during gene set enrichment analysis. BMC Bioinformatics 8, 332 (2007).

34. M. Haudiquet, A. Buffet, O. Rendueles, E. P. C. Rocha, Interplay between the cell envelope and mobile genetic elements shapes gene flow in populations of the nosocomial pathogen Klebsiella pneumoniae. PLOS Biology 19, e3001276 (2021).

35. B. D. Davis, Nonfiltrability of the agents of genetic recombination in Escherichia coli. J Bacteriol 60, 507–508 (1950).

36. A. L. Samuels, E. Lanka, J. E. Davies, Conjugative junctions in RP4-mediated mating of Escherichia coli. J Bacteriol 182, 2709–2715 (2000).

37. K. Moriguchi et al., Targeting Antibiotic Resistance Genes Is a Better Approach to Block Acquisition of Antibiotic Resistance Than Blocking Conjugal Transfer by Recipient Cells: A Genome-Wide Screening in Escherichia coli. Front Microbiol 10, 2939 (2019).

38. A. K. Vadakkepat et al., Cryo-EM structure of the R388 plasmid conjugative pilus reveals a helical polymer characterized by an unusual pilin/phospholipid binary complex. Structure 10.1016/j.str.2024.06.009 (2024).

39. S. E. George et al., Phenotypic heterogeneity and temporal expression of the capsular polysaccharide in Staphylococcus aureus. Molecular Microbiology 98, 1073–1088 (2015).

40. J. E. King, H. A. Aal Owaif, J. Jia, I. S. Roberts, Phenotypic Heterogeneity in Expression of the K1 Polysaccharide Capsule of Uropathogenic Escherichia coli and Downregulation of the Capsule Genes during Growth in Urine. Infect Immun 83, 2605–2613 (2015).

41. L. M. Reyes Ruiz, C. L. Williams, R. Tamayo, Enhancing bacterial survival through phenotypic heterogeneity. PLOS Pathogens 16, e1008439 (2020).

42. A. Nucci et al., Phenotypic heterogeneity of capsule production across opportunistic pathogens. mBio 0, e01807–01825 (2025).

43. K. A. Walker, V. L. Miller, The intersection of capsule gene expression, hypermucoviscosity and hypervirulence in Klebsiella pneumoniae. Current opinion in microbiology 54, 95–102 (2020).

44. M. A. Schembri, J. Blom, K. A. Krogfelt, P. Klemm, Capsule and fimbria interaction in Klebsiella pneumoniae. Infect Immun 73, 4626–4633 (2005).

45. D.-W. Wei et al., Insertion sequences accelerate genomic convergence of multidrug resistance and hypervirulence in Klebsiella pneumoniae via capsular phase variation. Genome Medicine 17, 45 (2025).

46. V. Cyriaque et al., Single-cell RNA sequencing reveals plasmid constrains bacterial population heterogeneity and identifies a non-conjugating subpopulation. Nature Communications 15, 5853 (2024).

47. R. J. B. Erickson et al., Single-Cell Analysis Reveals that the Enterococcal Sex Pheromone Response Results in Expression of Full-Length Conjugation Operon Transcripts in All Induced Cells. J Bacteriol 202 (2020).

48. S. Zobel et al., Tn7-Based Device for Calibrated Heterologous Gene Expression in Pseudomonas putida. ACS Synthetic Biology 4, 1341–1351 (2015).

49. N. R. Mattatall, K. E. Sanderson, Salmonella typhimurium LT2 possesses three distinct 23S rRNA intervening sequences. Journal of Bacteriology 178, 2272–2278 (1996).

50. T. Seemann, Snippy: rapid haploid variant calling and core SNP phylogeny. GitHub. Available at: github. com/tseemann/snippy (2015).

51. T. A. Gray et al., Intercellular communication and conjugation are mediated by ESX secretion systems in mycobacteria. Science 354, 347–350 (2016).

52. Y. Liao, G. K. Smyth, W. Shi, The Subread aligner: fast, accurate and scalable read mapping by seed-and-vote. Nucleic Acids Res 41, e108 (2013).

53. Y. Liao, G. K. Smyth, W. Shi, featureCounts: an efficient general purpose program for assigning sequence reads to genomic features. Bioinformatics 30, 923–930 (2014).

54. A. Alexa, J. Rahnenführer, Gene set enrichment analysis with topGO. Bioconductor Improv 27, 776 (2009).

55. C. P. Cantalapiedra, A. Hernández-Plaza, I. Letunic, P. Bork, J. Huerta-Cepas, eggNOG-mapper v2: Functional Annotation, Orthology Assignments, and Domain Prediction at the Metagenomic Scale. Molecular Biology and Evolution 38, 5825–5829 (2021).

56. P. D. Karp et al., The EcoCyc Database (2023). EcoSal Plus 11, eesp-0002-2023 (2023).

57. H. Wickham, C. Sievert, ggplot2: elegant graphics for data analysis (springer New York, 2009), vol. 10.

58. S. Hammerschmidt, M. Rohde, “Electron Microscopy to Study the Fine Structure of the Pneumococcal Cell“ in Streptococcus pneumoniae: Methods and Protocols, F. Iovino, Ed. (Springer New York, New York, NY, 2019), 10.1007/978-1-4939-9199-0_2, pp. 13–33.

59. T. R. Smallman, G. C. Williams, M. Harper, J. D. Boyce, Genome-Wide Investigation of Pasteurella multocida Identifies the Stringent Response as a Negative Regulator of Hyaluronic Acid Capsule Production. Microbiology Spectrum 10, e00195–00122 (2022).

60. S. A. C. Jillian C Danne and Jillian C Danne, Rachel Templin, Joan Clark, Viola Oorschot, Georg Ramm, Preparation of cells for transmission electron microscopy ultrastructural analysis. protocols.io dx. 10.17504/protocols.io.j8nlk47ewg5r/v1 (2022).

61. T. Guess et al., Size Matters: Measurement of Capsule Diameter in Cryptococcus neoformans. JoVE doi:10.3791/57171, e57171 (2018).

62. B. M. Coffey, G. G. Anderson, Biofilm formation in the 96-well microtiter plate. Methods Mol Biol 1149, 631–641 (2014).

63. E. A. Palombo, K. Yusoff, V. A. Stanisich, V. Krishnapillai, N. S. Willetts, Cloning and genetic analysis of tra cistrons of the Tra 2/Tra 3 region of plasmid RP1. Plasmid 22, 59–69 (1989).

64. Y. Fu et al., Mechanisms of blaIMP-4 dissemination across diverse carbapenem-resistant clinical isolates. Journal of Global Antimicrobial Resistance 41, 189–194 (2025).

65. M. M. C. Lam et al., Population genomics of hypervirulent Klebsiella pneumoniae clonal-group 23 reveals early emergence and rapid global dissemination. Nature Communications 9, 2703 (2018).

66. I. R. Lee et al., Differential host susceptibility and bacterial virulence factors driving Klebsiella liver abscess in an ethnically diverse population. Scientific Reports 6, 29316 (2016).

67. A. C. Jacobs et al., AB5075, a Highly Virulent Isolate of Acinetobacter baumannii, as a Model Strain for the Evaluation of Pathogenesis and Antimicrobial Treatments. mBio 5, e01076–01014 (2014).

68. L. A. Gallagher et al., Resources for Genetic and Genomic Analysis of Emerging Pathogen Acinetobacter baumannii. J Bacteriol 197, 2027–2035 (2015).

69. R. F. Wang, S. R. Kushner, Construction of versatile low-copy-number vectors for cloning, sequencing and gene expression in Escherichia coli. Gene 100, 195–199 (1991).

70. B. Bartolomé, Y. Jubete, E. Martínez, F. de la Cruz, Construction and properties of a family of pACYC184-derived cloning vectors compatible with pBR322 and its derivatives. Gene 102, 75–78 (1991).

71. J. P. Fürste et al., Molecular cloning of the plasmid RP4 primase region in a multi-host-range tacP expression vector. Gene 48, 119–131 (1986).

